# 5-Hydroxymethylcytosine signatures in cell-free DNA provide information about tumor types and stages

**DOI:** 10.1101/162081

**Authors:** Chun-Xiao Song, Senlin Yin, Li Ma, Amanda Wheeler, Yu Chen, Yan Zhang, Bin Liu, Junjie Xiong, Weihan Zhang, Jiankun Hu, Zongguang Zhou, Biao Dong, Zhiqi Tian, Stefanie S. Jeffrey, Mei-Sze Chua, Samuel So, Weimin Li, Yuquan Wei, Jiajie Diao, Dan Xie, Stephen R. Quake

## Abstract

5-Hydroxymethylcytosine (5hmC) is an important mammalian DNA epigenetic modification that has been linked to gene regulation and cancer pathogenesis. Here we explored the diagnostic potential of 5hmC in circulating cell-free DNA (cfDNA) using a sensitive chemical labeling-based low-input shotgun sequencing approach. We sequenced cell-free 5hmC from 49 patients of seven different cancer types and found distinct features that could be used to predict cancer types and stages with high accuracy. Specifically, we discovered that lung cancer leads to a progressive global loss of 5hmC in cfDNA, whereas hepatocellular carcinoma and pancreatic cancer lead to disease-specific changes in the cell-free hydroxymethylome. Our proof of principle results suggest that cell-free 5hmC signatures may potentially be used not only to identify cancer types but also to track tumor stage in some cancers.

**One Sentence Summary:** Analyzing the epigenetic modification 5-hydroxymethylcysoine in circulating cell-free DNA reveals tumor tissue of origin and stages for cancer diagnostics.

## Introduction

Circulating cell-free DNA (cfDNA) are DNA fragments found in the blood that originate from cell death in different tissues; this phenomenon has formed the basis of noninvasive prenatal diagnostic tests (*1*), organ transplant rejection diagnostics (*2*), and cancer detection (*3*). Recent work has focused on the identification of 5-methylcytosine (5mC) modifications in cfDNA to characterize a variety of potential health conditions (*3-8*). However, there has been no investigation to date of alternative epigenetic DNA modifications in cfDNA, due in part to the challenges of identifying and sequencing alternate modifications in low input DNA samples.

5-Hydroxymethylcytosine (5hmC) is a recently identified epigenetic mark which impacts a broad range of biological processes ranging from development to pathogenesis (*9,* 10). 5hmC is generated from 5mC by the TET family dioxygenases (*11*). Compared to the repressive effect of 5mC, 5hmC is generally believed to have a permissive effect on gene expression (*12-15*). Unlike 5mC which is uniformly distributed among different tissues in terms of total mass, 5hmC displays a tissue-specific mass distribution (*16*, 17) and low levels of 5hmC are also frequently observed in many solid tumors compared to corresponding normal tissues (*18*). These characteristics suggest that 5hmC may have potential value in cancer diagnostics (*10*). However, in contrast to the intensive studies on cell-free 5mC, cell-free 5hmC has remained unexploited, partly due to the low levels of 5hmC in the human genome (10 to 100-fold less than 5mC) (*17*) and the lack of a sensitive low-input 5hmC DNA sequencing method that would work with the minute amounts of cfDNA available (typically only a few nanograms per ml of plasma). In this work, we developed a sensitive chemical labeling-based low-input whole-genome 5hmC sequencing method that allows rapid and reliable sequencing of 5hmC in cfDNA, and showed that cell-free 5hmC display distinct features in several types of cancer, which can potentially be used not only to identify cancer types but also to track tumor stage in some cancers.

## Results

### Development of cell-free 5hmC sequencing

We developed a low-input whole-genome cell-free 5hmC sequencing method based on selective chemical labeling (hMe-Seal) (*13*). hMe-Seal is a robust method that uses β-glucosyltransferase (βGT) to selectively label 5hmC with a biotin *via* an azide-modified glucose for pull-down of 5hmC-containing DNA fragments for sequencing (*13*) (fig. S1A). Standard hMe-Seal procedure requires micrograms of DNA. In our modified approach, cfDNA is first ligated with sequencing adapters and 5hmC is selectively labeled with a biotin group. After capturing cfDNA containing 5hmC using streptavidin beads, the final library is completed by PCR directly from the beads instead of eluting the captured DNA to minimize sample loss during purification steps (Fig. 1A). With this approach we can sequence cell-free 5hmC readily from 1-10 ng of cfDNA. By spiking in a pool of 180 bp amplicons bearing C, 5mC, or 5hmC to cfDNA, we demonstrated that only 5hmC-containing DNA can be detected by PCR from the beads after pull-down (fig. S1B). This result was confirmed in the final sequencing libraries, which showed over 100-fold enrichment in reads mapping to 5hmC spike-in DNA (Fig. 1B). Furthermore, our approach performed equally well with cfDNA and bulk genomic DNA (1 μg whole blood genomic DNA (gDNA)) (Fig. 1B). The final cell-free 5hmC libraries are highly complex with a median unique nonduplicate map rate of 0.75 when lightly sequenced (median 15 million reads, ~ 0.5-fold human genome coverage) (fig. S1, C and D, and table S1), and yet technical replicates are highly reproducible (fig. S1E). We identified 5hmC-enriched regions (hMRs) in the sequence data using a Poisson-based method (*19*). hMRs are highly concordant between technical replicates and a pooled sample: over 75% of hMRs in the pooled sample are in common with each of the replicates (fig. S1F), reaching the ENCODE standard for ChIP-Seq (*20*). These results demonstrate that cell-free 5hmC can be readily and reliably profiled by the modified hMe-Seal method.

**Fig. 1.**
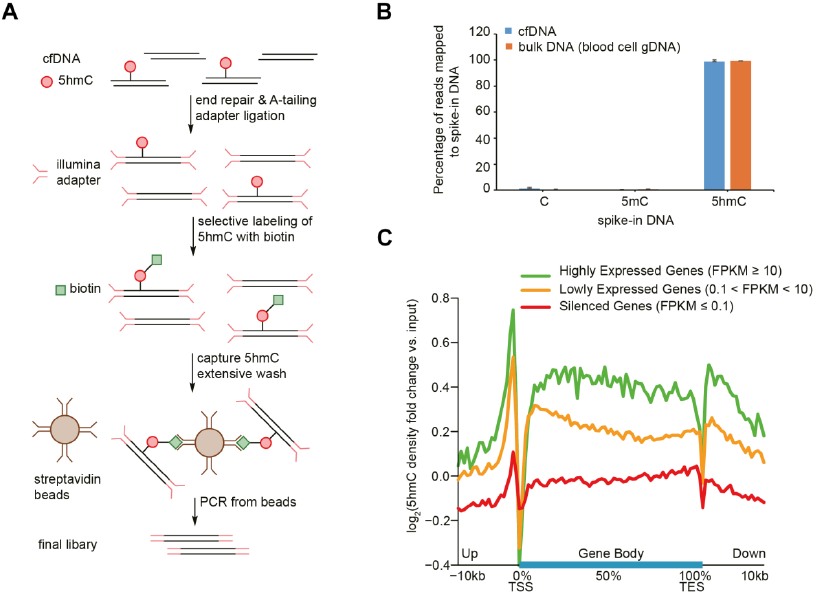
Sequencing of 5hmC in cfDNA. (**A**) General procedure of cell-free 5hmC sequencing. cfDNA is ligated with Illumina adapter and labeled with biotin on 5hmC for pull-down with streptavidin beads. The final library is completed by directly PCR from streptavidin beads. (**B**) Percentage of reads mapped to spike-in DNA in the sequencing libraries. Error bars indicate s.d. (**C**) Metagene profiles of log2 fold change of cell-free 5hmC to input cfDNA ratio in genes ranked according to their expression in cell-free RNA-Seq.

### Genome-wide mapping of 5hmC in cfDNA

We first sequenced cell-free 5hmC from eight healthy individuals (tables S1 and S2). We also sequenced 5hmC from whole blood gDNA from two of the individuals as blood is the major contributor to cell-free nucleic acids. Genome-scale profiles showed that the cell-free 5hmC distributions are nearly identical between healthy individuals and are clearly distinguishable from both the whole blood 5hmC distribution and the input cfDNA (fig. S2A). Previous studies of 5hmC in mouse and human tissues showed that the majority of 5hmC resides in the gene bodies and promoter proximal regions of the genome (*12*, 14). Genome-wide analysis of hMRs in our cfDNA data showed that a majority (80%) are intragenic with most enrichment in exons (observed to expected, o/e = 7.29), and depletion in intergenic regions (o/e = 0.46), consistent with that in whole blood (fig. S2, B and C) and in other tissues (*12,14*). The enrichment of 5hmC in gene bodies is known to be correlated with transcriptional activity in tissues such as the brain and liver (*12-14*). To determine whether this relationship holds in cfDNA, we performed sequencing of the cell-free RNA from the same individual (*21*). By dividing genes into three groups according to their cell-free RNA expression and plotting the average cell-free 5hmC profile alone gene bodies (metagene analysis), we discovered an enrichment of 5hmC in and around gene bodies of more highly expressed genes (Fig. 1C). These results demonstrate that cell-free 5hmC is derived from various tissue types and contains information from tissues other than the blood.

Since cell-free 5hmC were mostly enriched in the intragenic regions, we next used genic 5hmC fragments per kilobase of gene per million mapped reads (FPKM) to further compare the cell-free hydroxymethylome with the whole blood hydroxymethylome. Indeed, unbiased analysis of genic 5hmC using t-distributed stochastic neighbor embedding (tSNE) (*22*) showed strong separation between the cell-free and whole blood samples (fig. S2D). We used the *limma* package (*23*) to identify 2,082 differentially hydroxymethylated genes between whole blood and cell-free samples (*q*-values (Benjamini and Hochberg adjusted *p*-values) < 0.01, fold change > 2, fig. S3A). Notably, the 735 blood-specific 5hmC enriched genes showed increased expression in whole blood compared to the 1,347 cell-free-specific 5hmC enriched genes (*24*) (*p*-value < 2.2 × 10^-16^, Welch t-test) (fig. S3B). In agreement with the differential expression, Gene Ontology (GO) analysis (*25*) of blood-specific 5hmC enriched genes mainly identified blood cell-related processes (fig. S3C), whereas cell-free-specific 5hmC enriched genes identified much more diverse biological processes (fig. S3D). Examples of whole blood-specific (FPR1, FPR2) and cell-free-specific (GLP1R) 5hmC enriched genes are shown in fig. S3E. Together, these results provide further evidence that a variety of tissues contribute 5hmC to cfDNA and that measurement of this is a rough proxy for gene expression.

### Stage-dependent loss of 5hmC in lung cancer cfDNA

To explore the diagnostic potential of cell-free 5hmC, we applied our method to sequence cfDNA of a panel of 49 treatment-naïve primary cancer patients, including 15 lung cancer, 10 hepatocellular carcinoma (HCC), 7 pancreatic cancer, 4 glioblastoma (GBM), 5 gastric cancer, 4 colorectal cancer, 4 breast cancer patients (tables S3 to S9). These patients vary from early stage cancer to late stage metastatic cancer. In lung cancer, we observed a progressive global loss of 5hmC enrichment from early stage non-metastatic lung cancer to late stage metastatic lung cancer compared to healthy cfDNA, and it gradually resembled that of the unenriched input cfDNA (Fig. 2A). Unbiased gene body analysis using tSNE also showed a stage-dependent migration of the lung cancer profile from the healthy profile into one resembling the unenriched input cfDNA (fig. S4A). Notably, even the early stage lung cancer samples are highly separated from the healthy samples (fig. S4A). We further confirmed the global hypohydroxymethylome events using other metrics. First, most differential genes in metastatic lung cancer (*q*-values < 1e-7, 1,159 genes) showed stage-dependent depletion of 5hmC compared to healthy samples (Fig. 2B). Second, the metagene profile showed a stage-dependent depletion of gene body 5hmC signal and resemblance of the unenriched input cfDNA (fig. S4B). Third, there is a dramatic decrease in the number of hMRs identified in lung cancer, especially in metastatic lung cancer compared to healthy and other cancer samples (Fig. 2C). These data collectively indicate stage-dependent global loss of 5hmC levels in lung cancer cfDNA.

**Fig. 2.**
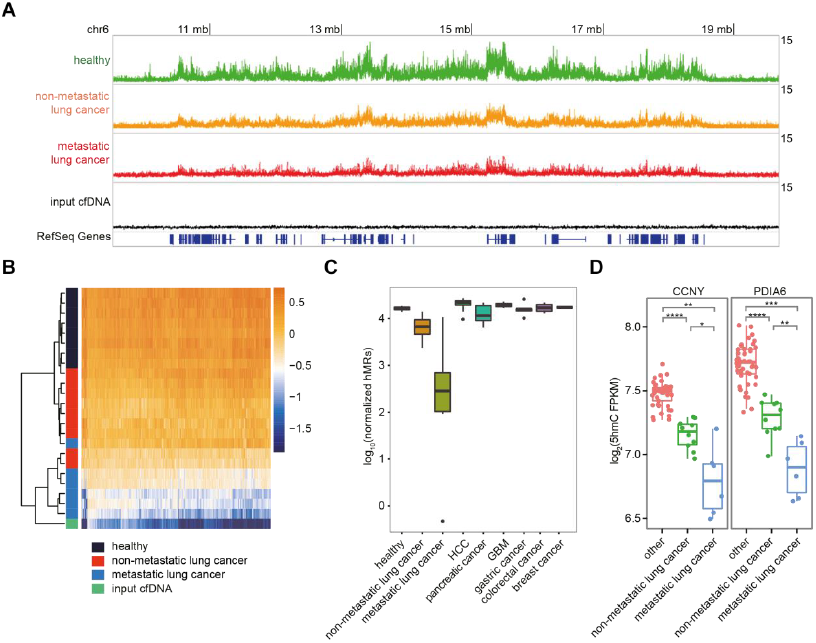
Lung cancer leads to progressive loss of 5hmC enrichment in cfDNA. (**A**) Genome browser view of the cell-free 5hmC distribution in a 10 mb region in chromosome 6. Showing the overlap tracks of healthy, non-metastatic lung cancer, metastatic lung cancer, and input cfDNA samples in line plot. (**B**) Heatmap of 1,159 metastatic lung cancer differential genes in healthy, lung cancer samples and the unenriched input cfDNA. Hierarchical clustering was performed across genes and samples. (**C**) Boxplot of number of hMRs (normalized to 1 million reads) identified in each group. (**D**) Boxplots of CCNY and PDIA6 5hmC FPKM in lung cancer and other cfDNA samples. \**P* < 0.05, \**P* <0.01,*** *P*<0.001,**** *P* <1e-5, Welch t-test.

It should be noted that the global loss of 5hmC enrichment seen in lung cancer cfDNA is not due to the failure of our enrichment method, as the spike-in control in all samples including the lung cancer samples showed high enrichment of 5hmC-containing DNA (fig. S4C). It is also a phenomenon unique to lung cancer that is not observed in other cancers we tested, evidenced by the number of hMRs (Fig. 2C) and the metagene profiles (fig. S4B). Examples of 5hmC depleted genes in lung cancer are shown in Fig. 2D and fig. S4D. Lung cancer tissue is known to have a low level of 5hmC compared to normal lung tissue (*18*), and lung has a relatively large contribution to cfDNA (*21*). It is plausible that lung cancer, especially metastatic lung cancer, causes large quantities of hypohydroxymethylated gDNA to be released into cfDNA, effectively diluting the cfDNA and leading to the depletion of 5hmC in the cell-free 5hmC landscape. Alternatively or in combination, the cfDNA hypohydroxymethylation could originate from blood gDNA hypohydroxymethylation observed in metastatic lung cancer patients as recently reported (*26*). Taken together these results indicate that cell-free 5hmC sequencing may potentially serve as a powerful tool for early lung cancer detection as well as monitoring lung cancer progression and metastasis.

### Monitoring treatment and recurrence in HCC

For HCC, we also sequenced cell-free 5hmC from seven patients with hepatitis B (HBV) infection, since most HCC cases are secondary to viral hepatitis infections (table S4). Unbiased gene level analysis by tSNE revealed that there is a gradual change of cell-free 5hmC from healthy to HBV and then to HCC, mirroring the disease development (Fig. 3A). HCC-specific differential genes (*q*-values < 0.001, fold change > 1.41, 1,006 genes) could separate HCC from healthy and most of the HBV samples (Fig. 3B). Both HCC-specific enriched and depleted genes can be identified compared to other cfDNA samples (Fig. 3B), and the enriched genes (379 genes) showed increased expression in liver tissue compared to the depleted genes (637 genes) (*24*) (*p*-values < 2.2 × 10^-16^, Welch t-test) (Fig. S5A), consistent with the permissive effect of 5hmC on gene expression. An example of HCC-specific 5hmC enriched genes is AHSG, a secreted protein highly expressed in the liver (*24*) (Fig. 3C and fig. S5, B and C), and an example of 5hmC depleted genes is TET2, one of enzyme that generate 5hmC and a tumor suppressor downregulated in HCC (*27*) (Fig. 3D and fig. S5D). Together, these results point to a model where virus infection and HCC development lead to a gradual damage of liver tissue and increased presentation of liver DNA in the blood.

**Fig. 3.**
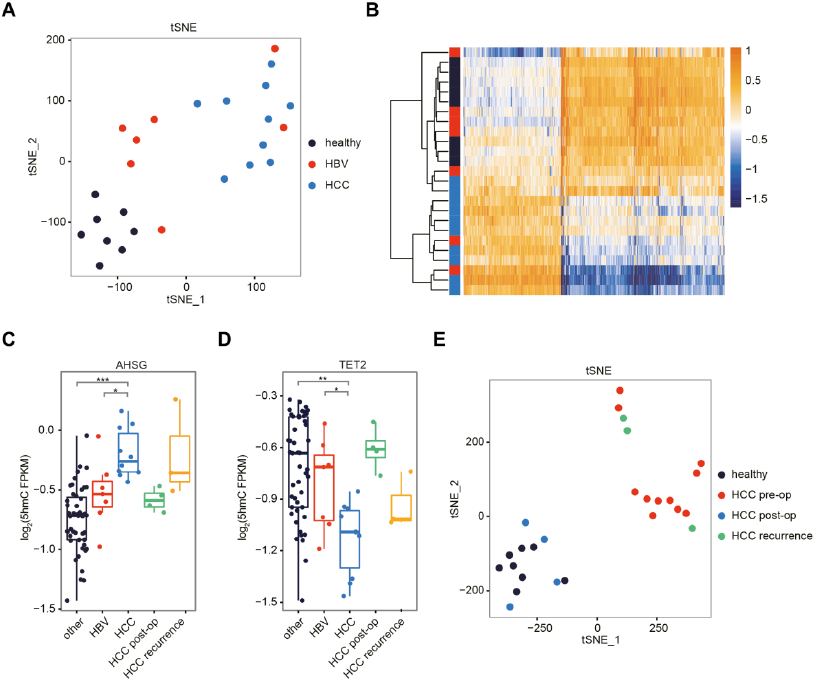
Cell-free 5hmC for monitoring HCC progression and treatment. (**A**) tSNE plot of 5hmC FPKM from healthy, HBV and HCC samples. (**B**) Heatmap of 1,006 HCC differential genes in healthy, HBV and HCC samples. Hierarchical clustering was performed across genes and samples. (**C** to **D**) Boxplots of AHSG (**C**) and TET2 (**D**) 5hmC FPKM in HBV, HCC (pre-op), HCC post-op, HCC recurrence and other cfDNA samples. \**P* < 0.05, \*\**P*<1e-4, \*\*\**P* <1e-5, Welch t-test. (**E**) tSNE plot of 5hmC FPKM from healthy, HCC pre-op, HCC post-op and HCC recurrence samples.

To further explore the potential of cell-free 5hmC for monitoring treatment and disease progression, we followed four of the HCC patients who underwent surgical resection, out of which three of them had recurrent disease (table S4). Analysis of serial plasma samples from these patients (pre-operation/pre-op; post-operation/post-op; and recurrence) with tSNE revealed that post-op samples clustered with healthy samples, whereas the recurrence samples clustered with HCC (Fig. 3E). This pattern was also reflected by changes in the 5hmC FPKM of AHSG and TET2 (Fig. 3, C and D). As an example of using cell-free 5hmC for tracking HCC treatment and progression, we employed linear discriminant analysis (LDA) to define a linear combination of the HCC-specific differential genes (Fig. 3D) into to a single value (the HCC score) that best separated the pre-op HCC samples from the healthy and HBV samples. We then calculated the HCC score for the post-op and recurrence HCC samples, and showed that the HCC score can accurately track the treatment and recurrence states (fig. S5E). Together, these results indicate that cell-free 5hmC sequencing presents an opportunity to detect HCC, as well as monitor treatment outcome and disease recurrence.

### Pancreatic cancer impacts the cell-free 5hmC

We also found pancreatic cancer produced drastic changes in its cell-free hydroxymethylome, even in some early stage pancreatic cancer patients (table S5). Like HCC, pancreatic cancer lead to both upregulated and downregulated 5hmC genes compared to healthy individuals (*q*-value < 0.01, fold change > 2, 713 genes) (fig. S6A). Examples of pancreatic cancer-specific 5hmC enriched and depleted genes compared other cfDNA samples are shown in fig. S6, B to E. Our results suggest that cell-free 5hmC sequencing can be potentially valuable for early detection of pancreatic cancer.

### Copy number variation (CNV) estimation

CNV can be detected from cfDNA sequencing, mostly in advanced cancer patients, which provides a way to assess the tumor burden in the cfDNA (*3*). To assess the tumor burden in our samples and to explore the relation between CNV contained from unenriched input cfDNA sequencing and the 5hmC enrichment sequencing, we also sequenced the input cfDNA in 47 samples (table S10). We analyzed the CNV from these input cfDNA sequencing with 1 mb bin (*28*), and as expected we can detect large scale CNV from about 20% of the cancer samples, mostly in late stage cancer samples (fig. S7A). We then analyzed the CNV from the corresponding 5hmC enrichment sequencing and interestingly, we found matched CNV patterns in several cases (fig. S7A). For example we could detect chromosome wise CNV in lung293 and lung417, two metastatic lung cancer samples, from input cfDNA sequencing (fig. S7, B and C). These samples display large scale cell-free 5hmC changes and correspondingly, the CNV patterns detected from the 5hmC enrichment sequencing mimic the CNV patterns detected from input cfDNA sequencing (fig. S7, D and E). This result supports the notion that 5hmC enriched cfDNA contains significant portion of tumor-derived cfDNA and therefore represents 5hmC patterns in tumor cells. It also shows that 5hmC sequencing and CNV analysis could complement each other in circulating tumor DNA analysis.

### Cancer type and stage prediction

Although there has been great interest in using cfDNA as a “liquid biopsy” for cancer detection, it has been challenging to identify the origin of tumor cfDNA and hence the location of the tumor. We discovered from tSNE analysis of all seven cancer types that lung cancer, HCC, and pancreatic cancer showed distinct signatures and could be readily separated from each other and healthy samples (Fig. 4A). The other four types of cancer displayed relatively minor changes compared to the healthy samples. Using other features such as the promotor region (5 kb upstream of the transcription start site (TSS)) showed similar patterns (fig. S8A). We note that no particular cancer type we tested resembled the whole blood profile (fig. S8B), suggesting that the blood cell contamination is not a significant source of variation. All patients in our panel fall in the same age range as the healthy individuals (fig. S8C, and tables S2 to S9), therefore age is unlikely to be a confounding factor. We also did not observe any batch effect (fig. S8D).

**Fig. 4.**
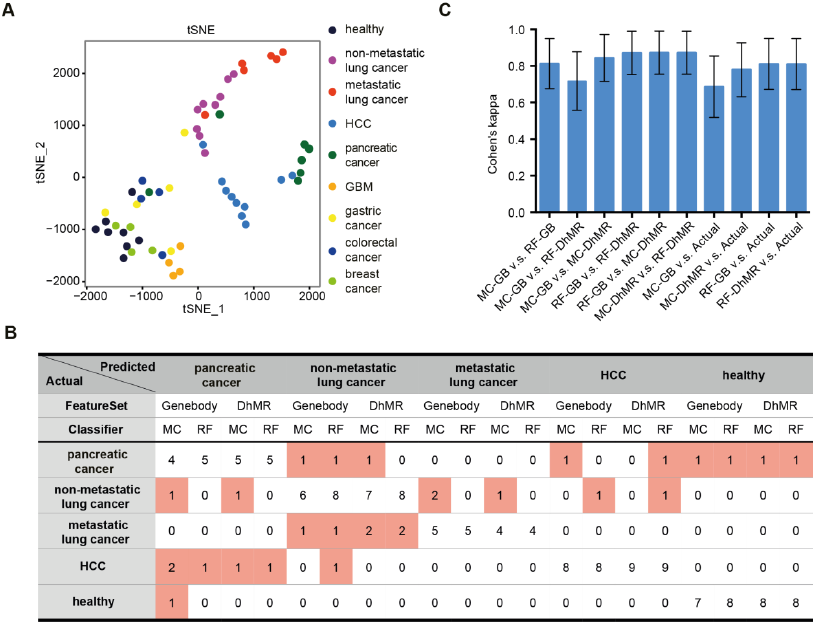
Cancer type and stage prediction with cell-free 5hmC. (**A**) tSNE plot of 5hmC FPKM in cfDNA from healthy and various cancer samples. (**B**) The actual and predicted classification by leave-one-out cross-validation using Mclust (MC) and Random Forest (RF) algorithm, based on two feature sets (gene body and DhMR). (**C**) The Cohen’s kappa coefficient for measuring inter-classifier agreement (GB for gene body). The error bar indicates 95% confidence interval of the Cohen’s kappa estimate.

To further demonstrate the potential of cfDNA 5hmC as a biomarker to predict cancer types we employed two widely used machine learning methods, the Gaussian mixture model (*29*) and Random Forest (*30*). We focused on the prediction HCC, pancreatic cancer, non-metastatic and metastatic lung cancer. Based on three rules (see Materials and Methods), we identified genes (table S11) whose average gene body 5hmC levels could either distinguish cancer groups from healthy groups or between cancer groups. In addition to using gene body data, the 5hmC on non-coding regions could also potentially serve as biomarkers in predicting cancer types (*9*). We therefore designed another set of features by investigating each of the 2kb windows of the entire genome and identified differential hMRs (DhMRs) for each cancer type (see Materials and Methods, and table S12). We trained the two machine learning algorithms using either differential 5hmC genes or DhMRs as features and evaluated the leave-one-out (LOO) cross-validation prediction accuracy. The Gaussian mixture model based predictor (Mclust) had overall successful prediction rates of 75% and 82.5%, when using gene body and DhMRs as features, respectively (Fig. 4B and fig. S9, A and B). Mclust-based dimensional reduction showed clear boundaries between the groups (fig. S9C). When only the type of the cancer is considered, Mclust predictors had higher success rate of 82.5% and 90% when using these two feature sets. The Random Forest predictor achieved LOO cross-validation prediction accuracy of 85%, when using either gene body or DhMRs as features (Fig. 4B). When only cancer type is considered, Random Forest predictor achieved 87.5% and 90% prediction accuracy, with gene body and DhMRs as features, respectively. Distinct 5hmC profiles in different cancer types of several DhMRs with high variable importance to random forest prediction model could be observed (fig. S9, D to E, and fig. S10). Finally, we used Cohen’s kappa to evaluate the concordance rate between different prediction models (*31*). All combinations showed high agreement (Cohen’s kappa ~ 0.8) in inter-classifier comparison and when comparing with the actual classification (Fig. 4C). These results support the prospects of using cell-free 5hmC for cancer diagnostics and staging.

## Discussion

Recent studies have reported that 5hmC is an important component of the mammalian genome (*9,32*). In this study, we reported an improved hMe-Seal (*13*) approach to sequence the low levels of 5hmC in cfDNA, which offers several notable advantages. First, unlike traditional bisulfite sequencing used for cell-free 5mC sequencing, our method does not further degrade the highly fragmented cfDNA. Second, compared to whole genome approaches including mutational sequencing and bisulfite sequencing, the enrichment for 5hmC not only enabling cost-effective sequencing (10-20 million reads, ~0.5-fold human genome coverage), but more importantly allowing the low frequency tissue contribution of 5hmC in cfDNA to be amplified from the dominant blood cell contribution in cfDNA.

We sequenced cell-free 5hmC from a panel of seven cancer types and focused our analysis on lung cancer, HCC and pancreatic cancer, the three cancers which displayed the most dramatic impact on the cell-free hydroxymethylome, even in the early stages. Lung and liver are reported to have relatively large contribution to cfDNA (*5,* 21), and pancreatic cancer is known to invade progressively to the lymph nodes and liver during early stages without remarkable symptoms, which may explain their large impact on the cell-free hydroxymethylome. In lung cancer we observed a characteristic stage-dependent global loss of cell-free 5hmC enrichment, while in HCC and pancreatic cancer, we identified significant finer scale changes of cell-free 5hmC (i.e. gene body and DhMR). In HCC, we also conducted an exploratory study of longitudinal samples whose results suggest that cell-free 5hmC may be used to monitor treatment and recurrence. Further studies will help elucidate how each cancer causes specific changes in the cell-free hydroxymethylome. Importantly, these three types of cancer displayed distinct patterns in their cell-free hydroxmethylome and we could employ machine learning algorithms trained with cell-free 5hmC features to predict the three cancer types with high accuracy.

In summary, we report the first proof-of-principle global analysis of hydroxymethylome in cfDNA. Large-scale clinical trials are required to fully validate the usefulness and understand potential limitations of this approach. Cell-free 5hmC contributes a new dimension of information to liquid biopsy-based diagnosis and prognosis; and we anticipate it may become a valuable tool for cancer diagnostics, as well as potentially for other disease areas, including but not limited to neurodegenerative diseases, cardiovascular diseases and diabetes. We envisage this strategy could be readily combined with other genetic and epigenetic-based cfDNA approaches (e.g. CNV analysis as we demonstrated) for increased diagnostic power. Our method represents the first enrichment-based genome-wide approach applied to cfDNA. The general framework of this method can be readily adopted to sequence other modifications in cell-free nucleic acids by applying the appropriate labeling chemistry to the modified bases. This would allow a comprehensive and global overview of genetic and epigenetic changes of various disease states, and further increase the power of personalized diagnostics.

## Materials and Methods

### Study design

The overall goal of this study was to explore the diagnostic potential of 5hmC cfDNA for cancer detection. The objective of the first portion of the study was to determine whether 5hmC can be sequenced from cfDNA using an enrichment-based method. The objective of the second portion of the study was to determine whether cell-free 5hmC contains information that can be used for cancer diagnostics. Samples for healthy subjects were obtained from Stanford blood center. HCC and breast cancer patients were recruited in a Stanford University Institutional Review Board-approved protocol. Lung cancer, pancreatic cancer, GBM, gastric cancer and colorectal cancer patients were recruited in a West China Hospital Institutional Review Board-approved protocol. All recruited subjects gave informed consent. No statistical methods were used to predetermine sample size. The experiments were not randomized and the investigators were not blinded to allocation during experiments and outcome assessment. No samples were excluded from the analysis.

### Clinical sample collection and processing

Blood was collected into EDTA-coated Vacutainers. Plasma was collected from the blood samples after centrifugation at 1,600 × *g* for 10 min at 4 °C and 16,000 × *g* at 10 min at 4 °C. cfDNA was extracted using the Circulating Nucleic Acid Kit (Qiagen). Whole blood genomic DNA was extracted using the DNA Mini Kit (Qiagen) and fragmented using dsDNA Fragmentase (NEB) into average 300 bp. DNA was quantified by Qubit Fluorometer (Life Technologies). Cell-free RNA was extracted using the Plasma/Serum Circulating and Exosomal RNA Purification Kit (Norgen). The extracted cell-free RNA was further digested using Baseline-ZERO DNases (Epicentre) and depleted using Ribo-Zero rRNA Removal Kit (Epicentre) according to a protocol from Clontech.

### Spike-in Amplicon Preparation

To generate the spiked-in control, lambda DNA was PCR amplified by Taq DNA Polymerase (NEB) and purified by AMPure XP beads (Beckman Coulter) in nonoverlapping ~180 bp amplicons, with a cocktail of dATP/dGTP/dTTP and one of the following: dCTP, dmCTP, or 10% dhmCTP (Zymo)/90% dCTP. Primers sequences are as follows: dCTP FW-CGTTTCCGTTCTTCTTCGTC, RV-TACTCGCACCGAAAATGTCA, dmCTP FW- GTGGCGGGTTATGATGAACT, RV-CATAAAATGCGGGGATTCAC, 10% dhmCTP/90% dCTP FW-TGAAAACGAAAGGGGATACG, RV-GTCCAGCTGGGAGTCGATAC.

### 5hmC Library Construction, Labeling, Capture and High-Throughput Sequencing

cfDNA (1-10 ng) or fragmented whole blood genomic DNA (1 μg) spiked with amplicons (0.01 pg of each amplicon per 10 ng DNA) was end repaired, 3’-adenylated and ligated to DNA Barcodes (Bioo Scientific) using KAPA Hyper Prep Kit (Kapa Biosystems) according to the manufacturer’s instructions. Ligated DNA was incubated in a 25 μL solution containing 50 mM HEPES buffer (pH 8), 25 mM MgCl_2_, 60 μM UDP-6-N_3_-Glc (Active Motif), and 12.5 U βGT (Thermo) for 2 hr at 37 °C. After that, 2.5 μL DBCO-PEG4-biotin (Click Chemistry Tools, 20 mM stock in DMSO) was directly added to the reaction mixture and incubated for 2 hr at 37 °C. Next, 10 μg sheared salmon sperm DNA (Life Technologies) was added into the reaction mixture and the DNA was purified by Micro Bio-Spin 30 Column (Bio–Rad). The purified DNA was incubated with 0.5 μL M270 Streptavidin beads (Life Technologies) pre-blocked with salmon sperm DNA in buffer 1 (5 mM Tris pH 7.5, 0.5 mM EDTA, 1 M NaCl and 0.2% Tween 20) for 30 min. The beads were subsequently undergone three 5-min washes each with buffer 1, buffer 2 (buffer 1 without NaCl), buffer 3 (buffer 1 with pH 9) and buffer 4 (buffer 3 without NaCl). All binding and washing were done at room temperature with gentle rotation. Beads were then resuspended in water and amplified with 14 (cfDNA) or 9 (whole blood genomic DNA) cycles of PCR amplification using Phusion DNA polymerase (NEB). The PCR products were purified using AMPure XP beads. Separate input libraries were made by direct PCR from ligated DNA without labeling and capture. For technical replicates, cfDNA from the same subject was divided into two technical replicates. Pair-end 75 bp sequencing was performed on the NextSeq instrument.

### Data Processing and Gene Body Analysis

FASTQ sequences were aligned to UCSC/hg19 with Bowtie2 v2.2.5 (*33*) and further filtered with samtools-0.1.19 (*34*) (view -f 2 -F 1548 -q 30 and rmdup) to retain unique non-duplicate matches to the genome. Pair-end reads were extended and converted into bedgraph format normalized to the total number of aligned reads using bedtools (*35*), and then converted to bigwig format using bedGraphToBigWig from the UCSC Genome Browser for visualization in Integrated Genomics Viewer (*36,* 37). FASTQ sequences were also aligned to the three spike-in control sequences to evaluate the pull-down efficiency. The spike-in control is only used as a validation of successful pull-down in each sample. hMRs were identified with MACS (*19*) using unenriched input DNA as background and default setting (*p*-value cutoff 1e-5). Genomic annotations of hMRs were performed by determining the percentage of hMRs overlapping each genomic regions ≥ 1 bp. Metagene profile was generated using ngs.plot (*38*). 5hmC FPKM were calculated using the fragment counts in each RefSeq gene body obtained by bedtools. For differential analyses, genes shorter than 1 kb or mapped to chromosome X and Y were excluded. Differential genic 5hmC analysis was performed using the *limma* package in R (*23*). GO analyses were performed using DAVID Bioinformatics Resources 6.7 with GOTERM_BP_FAT (*25,* 39). Tissue-specific gene expression was obtained from BioGPS (*24*, 40, 41). For tSNE plot, the Pearson correlation of gene body 5hmC FPKM was used as the distance matrix to tSNE. MA-plot, hierarchical clustering, tSNE, LDA, and heatmaps were done in R.

### Cell-free RNA Library Construction and High-Throughput Sequencing

Cell-free RNA library was prepared using ScriptSeq v2 RNA-Seq Library Preparation Kit (Epicentre) following the FFPE RNA protocol with 19 cycles of PCR amplification. The PCR products were then purified using AMPure XP beads. Pair-end 75 bp sequencing was performed on the NextSeq instrument. RNA-seq reads were first trimmed using Trimmomatic-0.33 (*42*) and then aligned using tophat-2.0.14 (*43*). RPKM expression values were extracted using cufflinks-2.2.1 (*44*) using RefSeq gene models.

### CNV estimation

The hg19 human genome was split into 1 mb bin and bin counts were generated using bedtools intersect. Mappability score of each bin was then assessed by average mappability using mappability track of hg19 from UCSC (kmer=75). Bins with mappability score under 0.8 were eliminated from further analysis. GC content percentages in each bin were calculated using getGC.hg19 from R package PopSV (1.0.0) and GC bias was corrected by fitting a LOESS model (correct.GC from R package PopSV 1.0.0). The corrected bin counts were then scaled by mean bin count of each sample and centered at 2. For estimation of CNV, we cutoff the corrected bin counts higher than 5 to minimize impact of extreme values. Moving averages with window size of 20 mb were then calculated within each chromosome, as the final estimation of CNV.

### Cancer type and stage prediction

Lung cancer, pancreatic cancer, HCC, and healthy samples (*n*=40) were included in the following analyses and leave-one-out (LOO) cross-validation was performed. With each iteration of LOO one sample was left out first, and the remaining 39 samples were used for feature selection and as a training dataset. The left out sample was then used to test the prediction accuracy of the machine learning model. Two types of feature selection were performed as independent analysis. In the “gene body” approach cancer type-specific marker genes were selected by performing a student t-test between 1) one cancer group and healthy group, 2) one cancer group and other cancer samples, 3) two different cancer groups. Benjamini and Hochberg correction was then performed for the raw *p*-value and the genes were then sorted by *q*-value. The top 5 genes with smallest *q*-value from each of these comparisons were selected as feature set to train the classifier. The second approach to finding features (“DhMR”) attempted to achieve higher resolution by first breaking the reference genome (hg19) into 2kb windows *in silico* and calculating 5hmC FPKM value for each of the window. Blacklisted genomic regions that tend to show artifact signal according to ENCODE were filtered before down-stream analysis (*45*). For cancer type-specific DhMRs, student t-test and Benjamini and Hochberg correction of *p*-values were performed for comparison pairs same as previously performed for identifying cancer-specific genes. The top 5 DhMRs with smallest *q*-value from each comparison were chosen for each cancer type. Random forest and Gaussian model-based Mclust classifier were performed on the dataset using previously described features (gene bodies and DhMRs). Classifiers were trained on lung cancer, pancreatic cancer, HCC and healthy samples. The same random seed (seed=5) was used in every random forest analysis for consistency. The top 15 features shared by at least 30 LOO iterations with the highest mean decrease Gini with the highest variable importance were plotted. Gaussian mixture model analysis was performed using Mclust R package (*29*). For Mclust model-based classifier training, a Bayesian information criterion (BIC) plot was performed for visualization of the classification efficacy of different multivariate mixture models. By default, the EEI model (diagonal, equal volume and shape) or VII model (spherical, unequal volume) with EDDA model-type (single component for each class with the same covariance structure among classes) were chosen for Mclust classification. Cohen’s kappa was then calculated for assessment of the interclassifier concordance.

### Statistical Analysis

We used unpaired two-tailed t-tests (Welch t-test) for normally distributed data in which two comparison groups were involved. In the case of multiple comparisons, Benjamini and Hochberg correction was then performed for the raw *p*-value to obtain the *q*-value. Random forest and Gaussian model-based Mclust were used as machine classifier. Cohen’s kappa was used for evaluating the predictive value of cell-free DNA 5hmC sequencing and interclassifier concordance. tSNE was used for dimension reduction and visualization. Statistical analyses were performed in R 3.3.2.

## Acknowledgements

We would like to acknowledge N. Neff and G. Mantalas for high-throughput sequencing; L. Penland and J. Beausang for sample collection; other members of the Quake and Xie labs for discussions and support; R. Altman and W. Zhou for critical reading of the manuscript.

## Author contributions

C.-X.S., J.D., D.X. and S.R.Q. conceived the study and designed the experiments. C.-X.S. performed the experiments with the help from L.M., Y.C., and B.D. C.-X.S. analyzed data with help from S.Y., Z.T. and D.X. L.M., A.W., Y.Z., B.L., J.X., W.Z.,J.H., Z.Z., S.S.J., M.-S.C., S.S., W.L., and Y.W. recruited patients, collected blood and organized clinical information. C.-X.S. and S.R.Q wrote the manuscript with input and comments from S.Y., D.X. and M.-S.C.

## Data and materials availability

All sequencing data were deposited in the Gene Expression Omnibus(http://www.ncbi.nlm.nih.gov/geo) under accession number GSE81314.

## Supplementary Materials

**Fig. S1.**
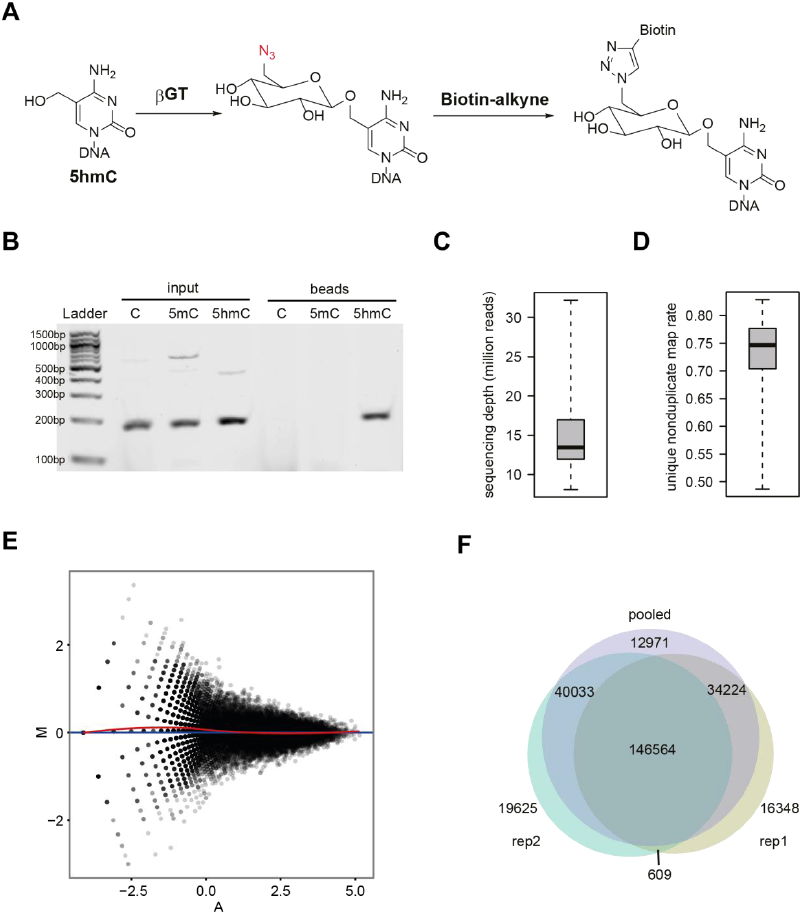
Cell-free 5hmC sequencing by modified hMe-Seal. (**A**) hMe-Seal reactions. 5hmC in DNA is labeled with an azide-modified glucose by βGT, which is then linked to a biotin group through click chemistry. (**B**) Enrichment tests of a single pool of amplicons containing C, 5mC or 5hmC spiked into cfDNA. Showing gel analysis that after hMe-Seal, only 5hmC-containing amplicon can be PCRed from the streptavidin beads. (**C**) Boxplot of sequencing depth across all cell-free samples. (**D**) Boxplot of unique nonduplicate map rate across all cell-free samples. (**E**) MA-plot of normalized cell-free 5hmC read counts (reads/million) in 10 kb bins genome-wide between technical duplicate. The horizontal blue line M = 0 indicates same value in two sample. A lowess fit (in red) is plotted underlying a possible trend in the bias related to the mean value. (**F**) Venn diagram of hMRs overlap between technical replications of cell-free 5hmC sequencing and a pooled sample from both replicates.

**Fig. S2.**
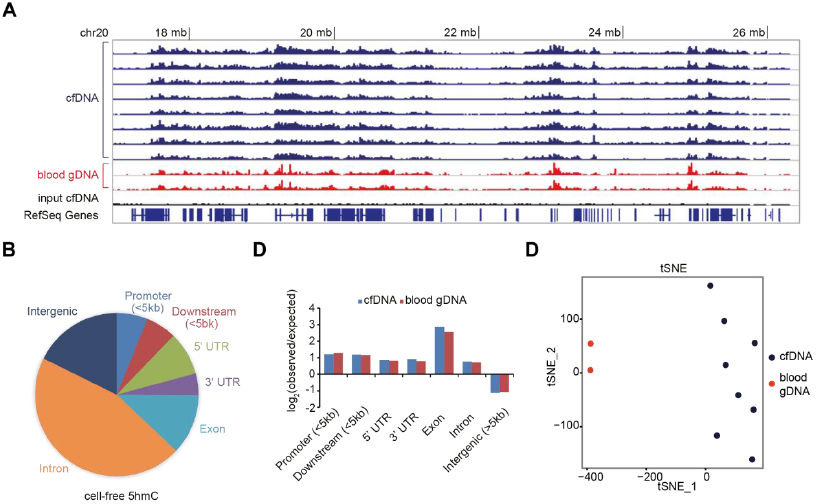
Genome-wide distribution of 5hmC in cfDNA. (**A**) Genome browser view of the 5hmC distribution in a 10 mb region in chromosome 20. Showing the tracks of enriched cfDNA and whole blood gDNA samples along with the unenriched input cfDNA. (**B**) Pie chart presentation of the overall genomic distribution of hMRs in cfDNA. (**C**) The relative enrichment of hMRs across distinct genomic regions in cfDNA and whole blood gDNA. (**D**) tSNE plot of 5hmC FPKM in cfDNA and whole blood gDNA from healthy samples.

**Fig. S3.**
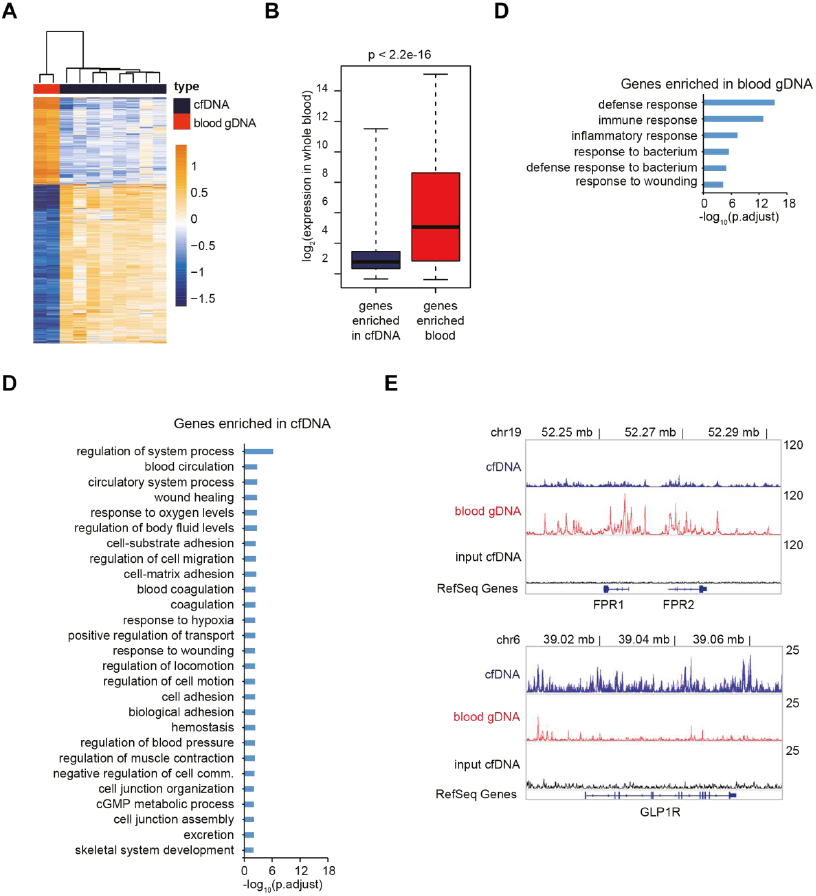
Differential 5hmC signals between cfDNA and whole blood gDNA. (**A**) Heatmap of 2,082 differential genes between cfDNA and blood gDNA. Hierarchical clustering was performed across genes and samples. (**B**) Boxplot of expression level in whole blood for cfDNA and whole blood gDNA 5hmC enriched genes. The *p*-value is shown on top. (**C** to **D**) GO analysis of the whole blood-specific (**C**) and cfDNA-specific (**D**) 5hmC enriched genes,adjusted *p*-value cut off 0.001. (**E**) Genome browser view of the 5hmC distribution in the FPR1/FPR2 (top) and the GLP1R (bottom) loci. Showing the overlap tracks of cfDNA, whole blood gDNA and input cfDNA in line plot.

**Fig. S4.**
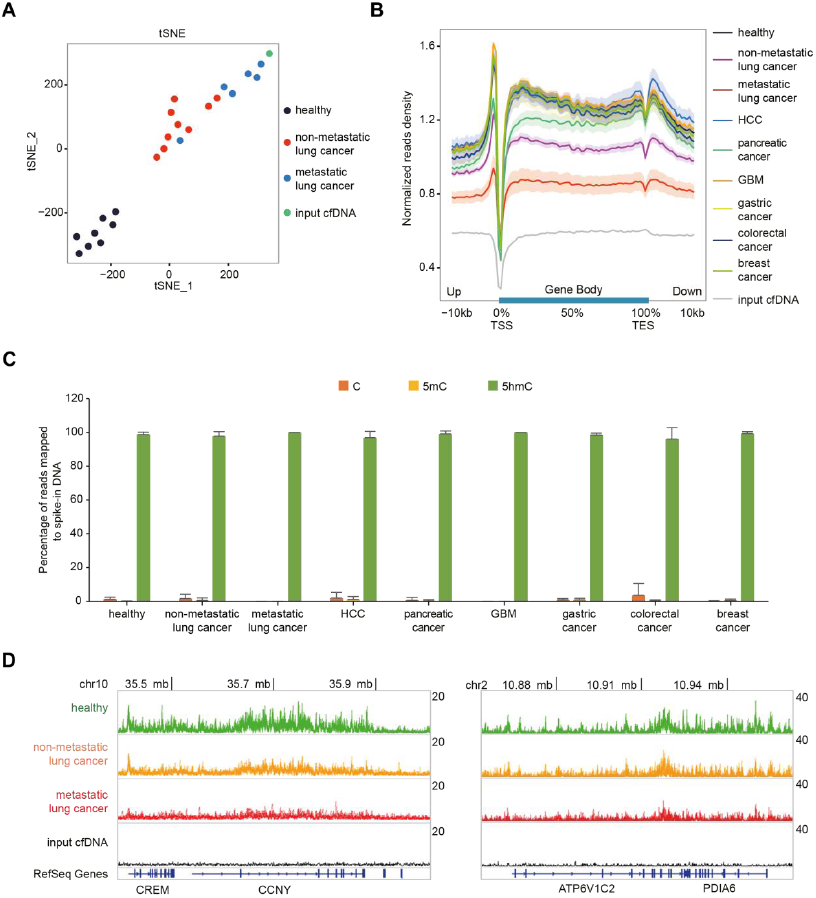
Cell-free hydroxymethylome in lung cancer. (**A**) tSNE plot of 5hmC FPKM from healthy, non-metastatic lung cancer and metastatic lung cancer samples, along with the unenriched input cfDNA. (**B**) Metagene profiles of cell-free 5hmC in healthy and various cancer groups, along with unenriched input cfDNA. Shaded area indicates s.e.m. (**C**) Percentage of reads mapped to spike-in DNA in the sequencing libraries of various groups.xError bars indicate s.d. (**D**) Genome browser view of the cell-free 5hmC distribution in the CREM/CCNY (left) and ATP6V1C2/PDIA6 (right) loci in healthy and lung cancer samples. Showing the overlap tracks in line plot.

**Fig. S5.**
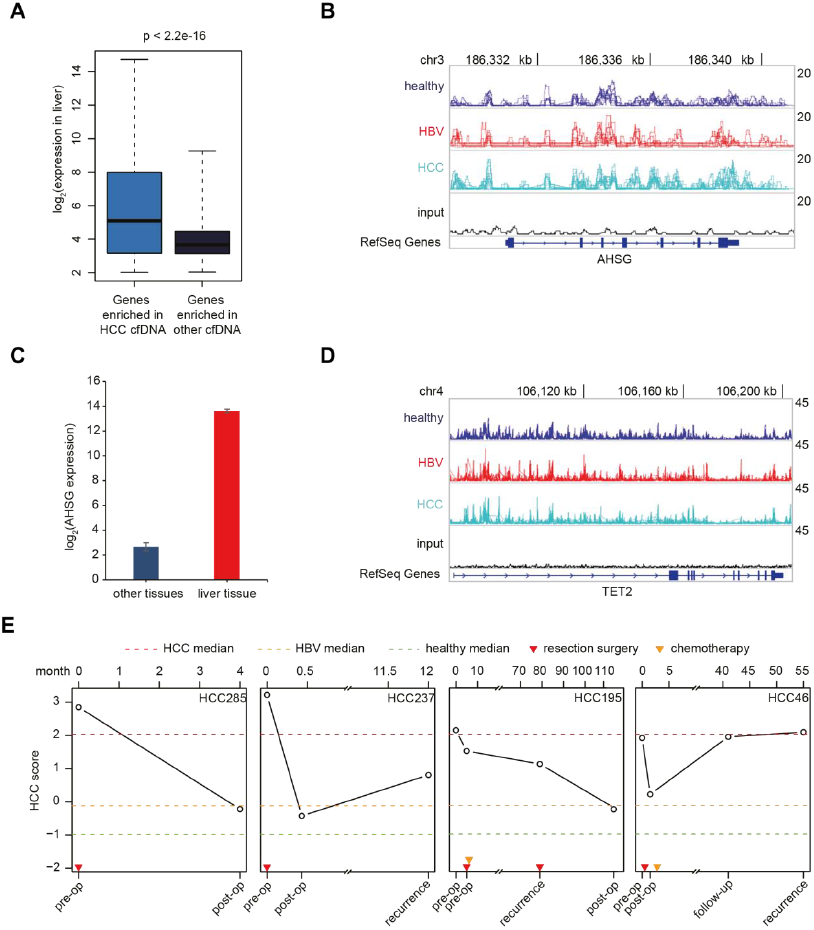
Cell-free hydroxymethylome in HCC. (**A**) Boxplot of expression level in liver tissue for HCC-specific 5hmC enriched and depleted genes. The *p*-value is shown on top. (**B**) Genome browser view of the cell-free 5hmC distribution in the AHSG locus in healthy HBV and HCC samples. Showing the overlap tracks in line plot. (**C**) Expression of AHSG in liver and other tissues. (**D**) Genome browser view of the cell-free 5hmC distribution in the TET2 locus in healthy, HBV and HCC samples. Showing the overlap tracks in line plot. (**E**) Changes of HCC score in 4 HCC follow-up cases. Disease status shown on the bottom. Time duration in month shown on the top. Dotted lines indicate the median values of HCC scores in the HCC, HBV, and healthy groups. Triangles indicate treatment. HCC score is a linear combination of 1,006 HCC differential genes (Fig. 3B) that best separates HCC from HBV and healthy samples.

**Fig. S6.**
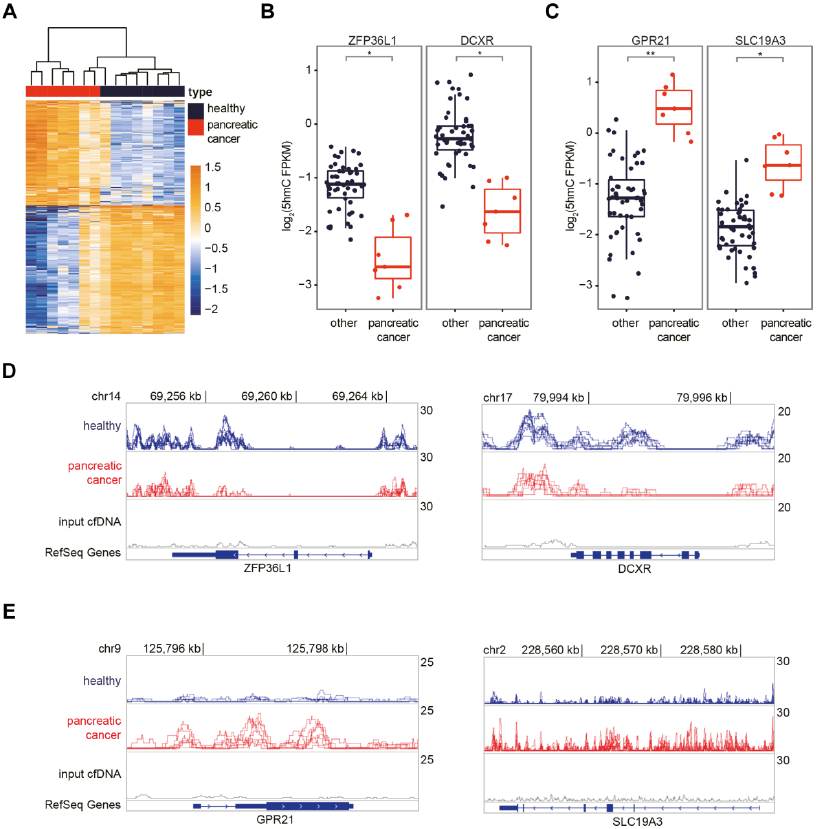
Cell-free hydroxymethylome in pancreatic cancer. (**A**) Heatmap of 713 pancreatic cancer differential genes in healthy and pancreatic cancer samples. Hierarchical clustering was performed across genes and samples. (**B** to **C**) Boxplots of ZFP36L1, DCXR (**B**) and GPR21, SLC19A3 (**C**) 5hmC FPKM in pancreatic cancer and other cfDNA samples. \**P* < 0.001, \*\**P* <1e-5, Welch t-test. (**D** to **E**) Genome browser view of the cell-free 5hmC distribution in the ZFP36L1, DCXR (**D**) and GPR21, SLC19A3 (**E**) loci in healthy and pancreatic cancer samples. Showing the overlap tracks in line plot.

**Fig. S7.**
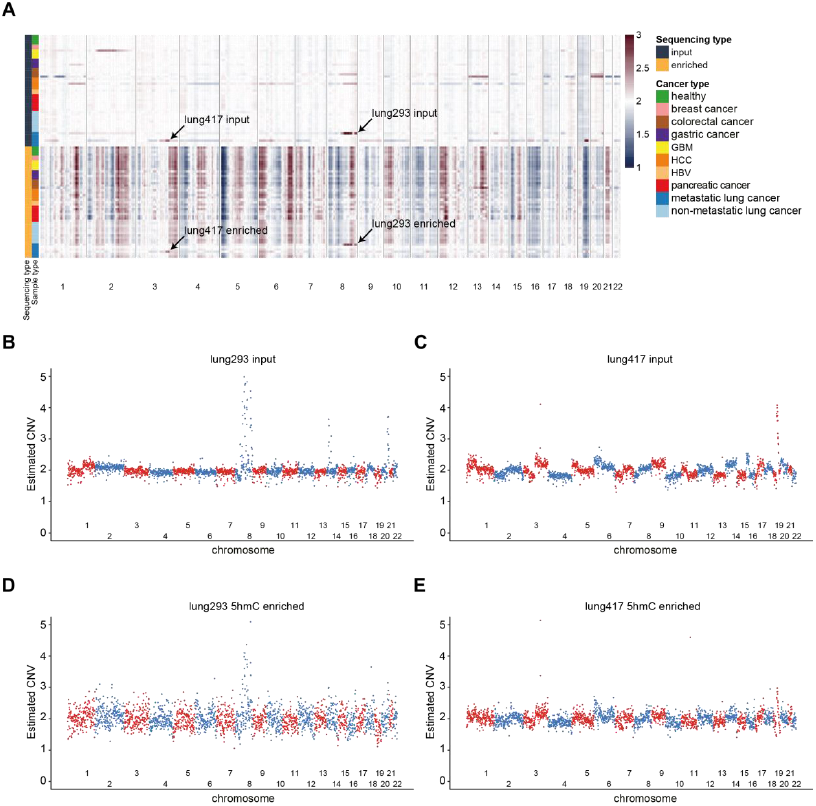
CNV estimation from input cfDNA and 5hmC enrichment sequencing. (**A**) CNV estimation heatmap from input cfDNA and 5hmC enrichment sequencing in 1 mb bin. Averaged bin counts within a sliding window of 20 bins were calculated as the estimated CNV for each bin. No clustering was performed. Arrows indicate samples with matched patterns in input cfDNA and 5hmC enrichment sequencing. (**B** to **C**) CNV estimation from input cfDNA sequencing of metastatic lung cancer patients lung293 (**B**) and lung417 (**C**). (**D** to **E**) CNV estimation from 5hmC enrichment sequencing of metastatic lung cancer patients lung293 (**D**) and lung417 (**E**).

**Fig. S8.**
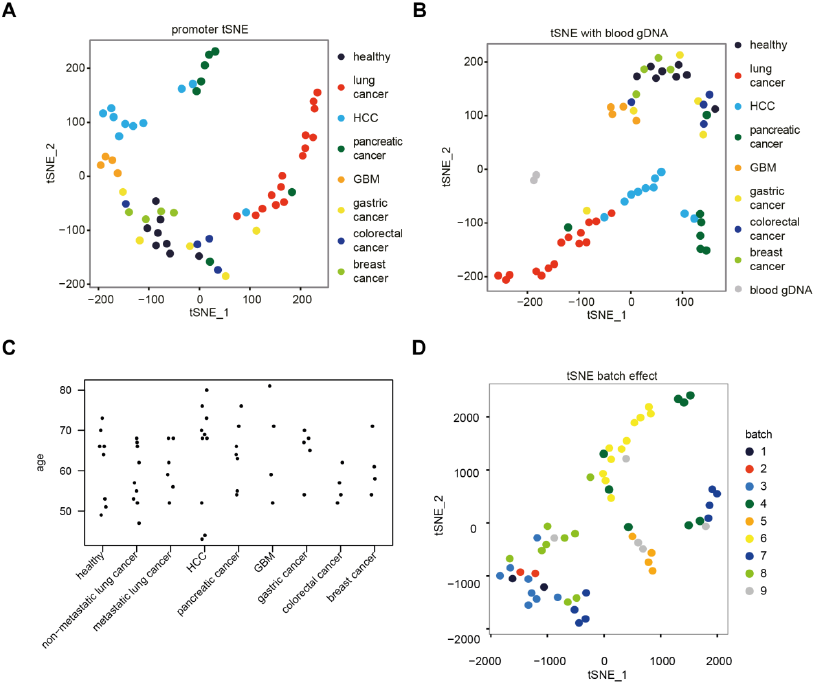
Cell-free hydroxymethylome in cancer samples. (**A**) tSNE plot of promoters 5hmC FPKM (5 kb upstream of TSS) from healthy and various caner samples. (**B**) tSNE plot of 5hmC FPKM from healthy and various caner cfDNA samples along with the whole blood gDNA samples. (**C**) Age distribution of healthy individual and various cancer patients. (**D**) tSNE plot of 5hmC FPKM in cfDNA from healthy and various cancer samples (Fig. 4A) colored by batches numbered according to the process time.

**Fig. S9.**
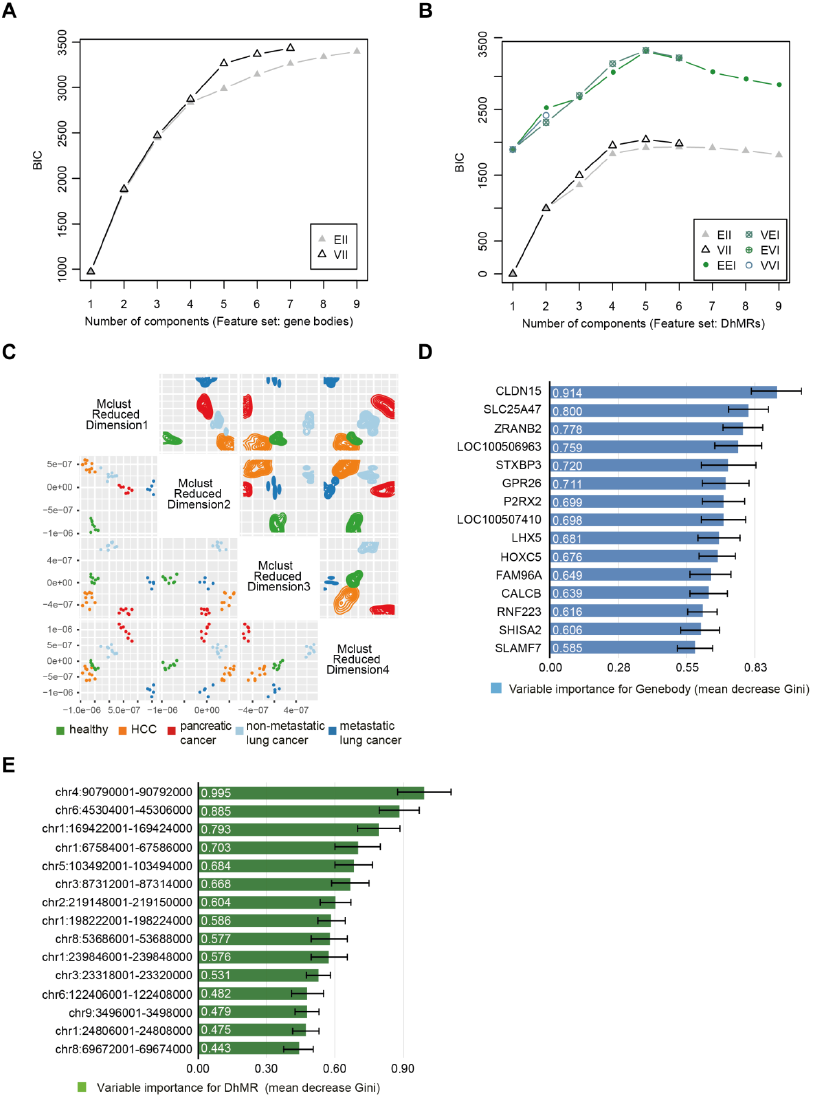
Cancer type and stage prediction with cell-free 5hmC. (**A** to **B**) Bayesian Information Criterion (BIC) plot by Mclust trained with 66 gene body feature set (**A**) and 71DhMRs feature set from samples other than lung324 (for leave-one-out cross-validation) (**B**), indicating high BIC value for separating five groups when using VII model for gene bodies and EEI for DhMRs. (**C**) 4-Dimensional Mclust-based dimensionality reduction plot using 66 gene body features from samples other than lung324 (for leave-one-out cross-validation). The lower half shows the scatter plot and the upper half shows the density plot. (**D** to **E**), Variable importance (mean decrease Gini) for the top 15 gene bodies (**D**) and DhMRs (**E**), in the random forest training model. The error bar indicates the standard deviation of variance importance from leave-one-out cross-validation.

**Fig. S10.**
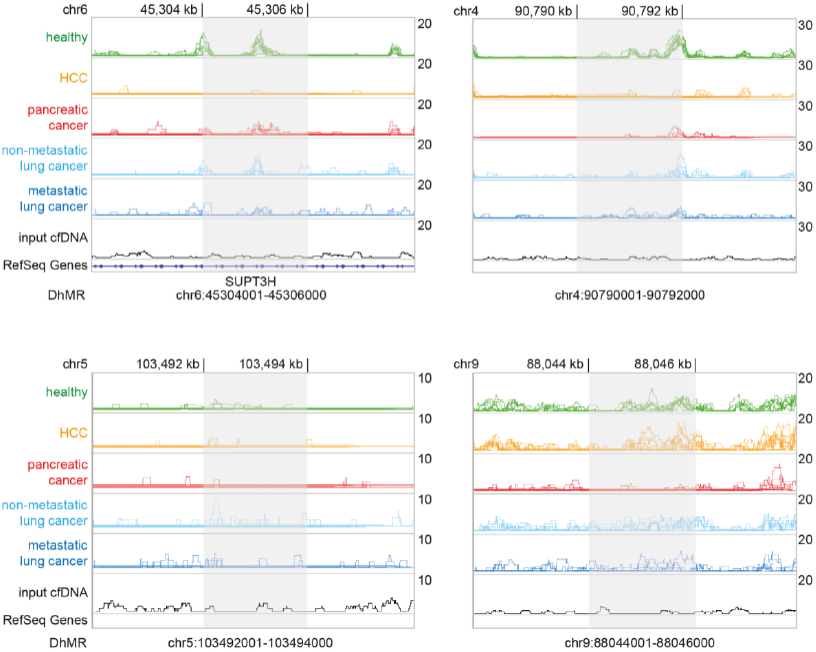
Examples of DhMRs in the random forest model. Genome browser view of the cell-free 5hmC distribution in four DhMRs with high variable importance in the random forest model (Fig. S9E) in various groups. Showing the overlap tracks in line plot. Shaded area indicates the DhMR.

## Supplemental Information

**Table S1.** Summary of 5hmC sequencing results.

| **sample ID** | **type** | **total reads**<br>**sequenced** | **unique nonduplicate**<br>**mapped reads** | **unique nonduplicate**<br>**mapped rate** |
| --- | --- | --- | --- | --- |
| **10** | healthy cfDNA | 20081973 | 15192613 | 0.76 |
| **11** | healthy cfDNA | 19142986 | 14762956 | 0.77 |
| **27** | healthy cfDNA | 21862078 | 16645192 | 0.76 |
| **35-1 \*** | healthy cfDNA | 29132339 | 16742468 | 0.57 |
| **35-2 \*** | healthy cfDNA | 28694218 | 17346511 | 0.60 |
| **36-1 \*** | healthy cfDNA | 32202519 | 20996955 | 0.65 |
| **36-2 \*** | healthy cfDNA | 31089686 | 20993595 | 0.68 |
| **38o** | healthy cfDNA | 20124203 | 15295376 | 0.76 |
| **38** | healthy cfDNA | 20419287 | 15679281 | 0.77 |
| **39o** | healthy cfDNA | 22320662 | 17833176 | 0.80 |
| **input §** | cfDNA input | 38574253 | 25910419 | 0.67 |
| **35-blood** | whole blood gDNA | 44077590 | 31654982 | 0.72 |
| **36-blood** | whole blood gDNA | 40843066 | 29266169 | 0.72 |
| **blood-input †** | whole blood gDNA input | 39138506 | 26455609 | 0.68 |
| **lung293** | lung cancer | 14172402 | 11470840 | 0.81 |
| **lung323** | lung cancer | 12269885 | 8916594 | 0.73 |
| **lung324** | lung cancer | 13313728 | 10058078 | 0.76 |
| **lung395** | lung cancer | 13589263 | 10092883 | 0.74 |
| **lung417** | lung cancer | 13212811 | 10109574 | 0.77 |
| **lung418** | lung cancer | 13103903 | 10420656 | 0.80 |
| **lung419** | lung cancer | 11949356 | 9704240 | 0.81 |
| **lung492** | lung cancer | 12563742 | 8885504 | 0.71 |
| **lung493** | lung cancer | 12930120 | 10479700 | 0.81 |
| **lung496** | lung cancer | 12267496 | 9657956 | 0.79 |
| **lung512** | lung cancer | 12934833 | 10483836 | 0.81 |
| **lung513** | lung cancer | 11310088 | 8304508 | 0.73 |
| **lung514** | lung cancer | 12895079 | 10264145 | 0.80 |
| **lung515** | lung cancer | 12132995 | 9406700 | 0.78 |
| **lung517** | lung cancer | 11766082 | 8857054 | 0.75 |
| **HCC150** | HCC | 15215190 | 11298385 | 0.74 |
| **HCC237** | HCC | 13439935 | 10109197 | 0.75 |
| **HCC241** | HCC | 16201676 | 12017320 | 0.74 |
| **HCC256** | HCC | 14579945 | 10728759 | 0.74 |
| **HCC260** | HCC | 13791503 | 10021911 | 0.73 |
| **HCC285** | HCC | 11522024 | 7662330 | 0.67 |
| **HCC290** | HCC | 13162465 | 9271065 | 0.70 |
| **HCC320** | HCC | 13462633 | 9696240 | 0.72 |
| **HCC341** | HCC | 11199473 | 6497400 | 0.58 |
| **HCC628** | HCC | 15365745 | 11759122 | 0.77 |
| **HCC324** | HCC | 12525818 | 9598812 | 0.77 |
| **HCC46** | HCC | 13121530 | 9237102 | 0.70 |
| **HCC73** | HCC | 13816686 | 10745247 | 0.78 |
| **HCC398** | HCC | 13791599 | 10430016 | 0.76 |
| **HCC489** | HCC | 11446887 | 5575387 | 0.49 |
| **HCC195** | HCC | 11538777 | 7701351 | 0.67 |
| **HCC234** | HCC | 11960087 | 8468478 | 0.71 |
| **HCC626** | HCC | 13552712 | 11087605 | 0.82 |
| **HCC647** | HCC | 12491614 | 8590321 | 0.69 |
| **pancreatic27** | pancreatic cancer | 9717087 | 8019436 | 0.83 |
| **pancreatic68** | pancreatic cancer | 10457109 | 8374219 | 0.80 |
| **pancreatic69** | pancreatic cancer | 10838005 | 8940883 | 0.82 |
| **pancreatic75** | pancreatic cancer | 10197772 | 8452749 | 0.83 |
| **pancreatic9** | pancreatic cancer | 14601356 | 11245279 | 0.77 |
| **pancreatic15** | pancreatic cancer | 15240467 | 11923009 | 0.78 |
| **pancreatic22** | pancreatic cancer | 13439343 | 10356395 | 0.77 |
| **GBM57** | GBM | 8799132 | 6455359 | 0.73 |
| **GBM58** | GBM | 8874810 | 7253089 | 0.82 |
| **GBM66** | GBM | 9795211 | 8073651 | 0.82 |
| **GBM76** | GBM | 8103209 | 6165341 | 0.76 |
| **stomach1** | gastric cancer | 14282633 | 10365849 | 0.73 |
| **stomach2** | gastric cancer | 17825012 | 12938872 | 0.73 |
| **stomach3** | gastric cancer | 16979690 | 12894400 | 0.76 |
| **stomach4** | gastric cancer | 21192604 | 15675499 | 0.74 |
| **stomach8** | gastric cancer | 14070772 | 8321549 | 0.59 |
| **colon13** | colorectal cancer | 17352371 | 12517451 | 0.72 |
| **colon16** | colorectal cancer | 15470656 | 11210513 | 0.72 |
| **colon17** | colorectal cancer | 15101557 | 10590748 | 0.70 |
| **colon19** | colorectal cancer | 18441208 | 12503926 | 0.68 |
| **BR5-1 \*** | breast cancer | 17826666 | 13542700 | 0.76 |
| **BR5-2 \*** | breast cancer | 17746176 | 13004851 | 0.73 |
| **BR7-1 \*** | breast cancer | 16963664 | 13160842 | 0.78 |
| **BR7-2 \*** | breast cancer | 15495003 | 12100951 | 0.78 |
| **BR13** | breast cancer | 21382473 | 16015986 | 0.75 |
| **BR14** | breast cancer | 18668112 | 14613260 | 0.78 |
| **HBV268** | HBV | 8730571 | 5106519 | 0.58 |
| **HBV334** | HBV | 11838111 | 7848078 | 0.66 |
| **HBV374** | HBV | 14896634 | 11099981 | 0.75 |
| **HBV397** | HBV | 12127855 | 8416798 | 0.69 |
| **HBV455** | HBV | 12796382 | 9001735 | 0.70 |
| **HBV640** | HBV | 10040349 | 6062886 | 0.60 |
| **HBV646** | HBV | 9665264 | 5002160 | 0.52 |

\* Technical duplicate.
§ Unenriched input cfDNA, mixed from samples 35 and 36.
† Unenriched input whole blood gDNA, mixed from samples 35-blood and 36-blood.

**Table S2.** Clinical information for healthy samples.

| sample ID | gender | age |
| --- | --- | --- |
| **10** | female | 53 |
| **11** | female | 66 |
| **27** | female | 66 |
| **35** | male | 51 |
| **36** | male | 73 |
| **38o** | female | 70 |
| **38** | female | 64 |
| **39o** | female | 49 |

**Table S3.**
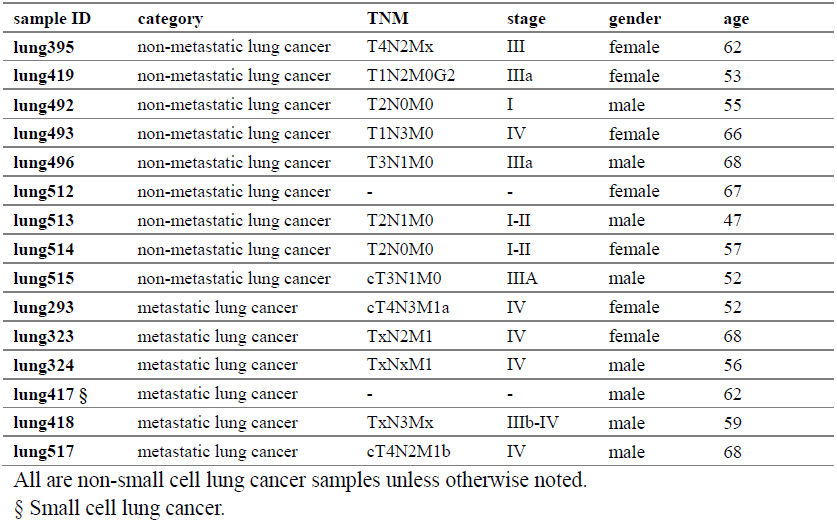
Clinical information for lung cancer samples.

**Table S4.** Clinical information for HCC samples.

| sample ID | category | TNM | tumor size (cm) | gender | age |
| --- | --- | --- | --- | --- | --- |
| **HBV268** | HBV | - | - | male | 36 |
| **HBV334** | HBV | - | - | female | 55 |
| **HBV374** | HBV | - | - | female | 45 |
| **HBV397** | HBV | - | - | female | 51 |
| **HBV455** | HBV | - | - | female | 66 |
| **HBV640** | HBV | - | - | female | 49 |
| **HBV646** | HBV | - | - | male | 60 |
| **HCC150** | HCC pre-op | pT1 pNX pMX | 3.1 § | male | 76 |
| **HCC256** | HCC pre-op | pT1 pNX pMX | 15x9 | male | 80 |
| **HCC260** | HCC pre-op | pT1 pNX pMX | 1.3 § | male | 68 |
| **HCC290** | HCC pre-op | - | 10x13x18 | male | 68 |
| **HCC320** | HCC pre-op | - | multifocal | female | 70 |
| **HCC628** | HCC pre-op | pT1 | 1.8 § | male | 43 |
| **HCC285** | HCC pre-op | pT3N0M0 | 8 § | male | 73 |
| **HCC324** | HCC post-op | - | - |  | 73 |
| **HCC237** | HCC pre-op | pT2 pNX pMX | 4.1 § | male | 52 |
| **HCC241** | HCC post-op | - | - |  | 52 |
| **HCC341** | HCC recurrence | - | 3x1.2 |  | 53 |
| **HCC195** | HCC pre-op | pT1 pNX pM0 | - | male | 44 |
| **HCC234** | HCC pre-op | - | 1.6 § |  | 44 |
| **HCC626** | HCC recurrence | pT1 pNX pM0 | 1.7x1.7x1.0 |  | 50 |
| **HCC647** | HCC post-op | - | - |  | 53 |
| **HCC46** | HCC pre-op | pT2 pNX pMX | 2.8 § | male | 69 |
| **HCC73** | HCC post-op | - | - |  | 69 |
| **HCC398** | HCC follow-up | - | - |  | 72 |
| **HCC489** | HCC recurrence | - | 2.2 § |  | 73 |

Same color shade indicate follow-up of the same patient.
§ in greatest dimension.

**Table S5.** Clinical information for pancreatic cancer samples.

| sample ID | TNM | stage | metastasis to | gender | age |
| --- | --- | --- | --- | --- | --- |
| **pancreatic9** | T3N0M1 | IV | liver | male | 76 |
| **pancreatic15** | T1N0M0 | IA | - | male | 64 |
| **pancreatic22** | T4N1M0 | III | - | female | 71 |
| **pancreatic27** | T4N1M1 | IV | abdominal wall, omentum | male | 55 |
| **pancreatic68** | T3N0M1 | IV | liver | male | 63 |
| **pancreatic69** | T3N0M0 | IIA | - | male | 66 |
| **pancreatic75** | T3N0M0 | IIA | - | male | 54 |

**Table S6.** Clinical information for GBM samples.

| sample ID | stage | gender | age |
| --- | --- | --- | --- |
| **GBM57** | IV | female | 52 |
| **GBM58** | IV | male | 71 |
| **GBM66** | IV | male | 81 |
| **GBM76** | IV | male | 59 |

**Table S7.** Clinical information for gastric cancer samples.

| sample ID | TNM | stage | gender | age |
| --- | --- | --- | --- | --- |
| stomach1 | T2N1M0 | II a | male | 67 |
| stomach2 | T4aN3bM0 | III c | male | 54 |
| stomach3 | T1aN0M0 | I a | male | 68 |
| stomach4 | T4bN0M0 | III b | male | 70 |
| stomach8 | T1bN0M0 | I a | male | 65 |

**Table S8.** Clinical information for colorectal cancer samples.

| sample ID | TNM | stage | gender | age |
| --- | --- | --- | --- | --- |
| colon13 | T4N0M0 | II | male | 54 |
| colon16 | T3N0M0 | II | female | 57 |
| colon17 | T4N0M1 | IV | male | 52 |
| colon19 | pT4N1M1 | IV | female | 62 |

**Table S9.** Clinical information for breast cancer samples.

| sample ID | tumor size (cm) | tumor grade | age |
| --- | --- | --- | --- |
| BR5 | 2.5 | 2 | 54 |
| BR7 | 1.2 | 1 | 71 |
| BR13 | 1 | 2 | 58 |
| BR14 | 1.9 | 1 | 61 |

**Table S10.** Summary of input cfDNA sequencing results.

| sample ID | type | total reads<br>sequenced | unique<br>nonduplicate<br>mapped reads | unique<br>nonduplicate<br>mapped rate |
| --- | --- | --- | --- | --- |
| 10-input | healthy | 12297454 | 9023854 | 0.73 |
| 27-input | healthy | 14185485 | 10553292 | 0.74 |
| 38o-input | healthy | 12534132 | 9265385 | 0.74 |
| 38-input | healthy | 12329856 | 9068759 | 0.74 |
| lung293-input | lung cancer | 14387649 | 10706785 | 0.74 |
| lung323-input | lung cancer | 12718973 | 9096119 | 0.72 |
| lung324-input | lung cancer | 14283775 | 10338094 | 0.72 |
| lung395-input | lung cancer | 12340896 | 8984866 | 0.73 |
| lung417-input | lung cancer | 17296343 | 12782946 | 0.74 |
| lung418-input | lung cancer | 20112071 | 13065287 | 0.65 |
| lung419-input | lung cancer | 17101222 | 12919260 | 0.76 |
| lung492-input | lung cancer | 13523477 | 8003249 | 0.59 |
| lung493-input | lung cancer | 14517791 | 10169937 | 0.70 |
| lung496-input | lung cancer | 11334027 | 8241433 | 0.73 |
| lung512-input | lung cancer | 14193573 | 10244580 | 0.72 |
| lung513-input | lung cancer | 17138456 | 10633764 | 0.62 |
| lung514-input | lung cancer | 14089241 | 9448179 | 0.67 |
| lung515-input | lung cancer | 12218453 | 8370255 | 0.69 |
| **lung517-input** | lung cancer | 123012581 | 74444363 | 0.61 |
| **HCC237-input** | HCC | 16868568 | 11576052 | 0.69 |
| **HCC241-input** | HCC | 19915649 | 13060838 | 0.66 |
| **HCC290-input** | HCC | 14729600 | 9148988 | 0.62 |
| **HCC628-input** | HCC | 13985304 | 9264263 | 0.66 |
| **HCC324-input** | HCC | 10706376 | 7158113 | 0.67 |
| **pancreatic27-input** | pancreatic cancer | 21005431 | 15092249 | 0.72 |
| **pancreatic68-input** | pancreatic cancer | 21609699 | 16086862 | 0.74 |
| **pancreatic69-input** | pancreatic cancer | 20381405 | 15078736 | 0.74 |
| **pancreatic75-input** | pancreatic cancer | 20150809 | 15023706 | 0.75 |
| **pancreatic9-input** | pancreatic cancer | 17788884 | 12676327 | 0.71 |
| **pancreatic15-input** | pancreatic cancer | 66823239 | 44834027 | 0.67 |
| **pancreatic22-input** | pancreatic cancer | 20343874 | 13994980 | 0.69 |
| **GBM57-input** | GBM | 16663028 | 12288601 | 0.74 |
| **GBM58-input** | GBM | 19745555 | 15066250 | 0.76 |
| **GBM66-input** | GBM | 22743166 | 16710159 | 0.73 |
| **GBM76-input** | GBM | 19426157 | 14503092 | 0.75 |
| **stomach1-input** | gastric cancer | 15593466 | 11708043 | 0.75 |
| **stomach2-input** | gastric cancer | 19726402 | 14537170 | 0.74 |
| **stomach4-input** | gastric cancer | 12241169 | 9077833 | 0.74 |
| **stomach8-input** | gastric cancer | 15604495 | 11004207 | 0.71 |
| **colon13-input** | colorectal cancer | 19419793 | 14360900 | 0.74 |
| **colon16-input** | colorectal cancer | 17016615 | 12384120 | 0.73 |
| **colon17-input** | colorectal cancer | 18873289 | 13685934 | 0.73 |
| **colon19-input** | colorectal cancer | 20046893 | 14638576 | 0.73 |
| **BR7-input** | breast cancer | 17555208 | 12726778 | 0.72 |
| **BR13-input** | breast cancer | 18015338 | 13467760 | 0.75 |
| **HBV397-input** | HBV | 12448351 | 8786039 | 0.71 |
| **HBV640-input** | HBV | 16143446 | 10547979 | 0.65 |

**Table S11.** Top gene body feature set used for cancer prediction.

|  |  |  |  |  |  |
| --- | --- | --- | --- | --- | --- |
| CLDN15 | SLC25A47 | ZRANB2 | LOC100506963 | STXBP3 | GPR26 |
| P2RX2 | LOC100507410 | LHX5 | HOXC5 | FAM96A | CALCB |
| RNF223 | SHISA2 | SLAMF7 | PAX1 | DACH1 | LOC100128946 |
| ASF1B | KIF16B | SSR2 | LARS | DHRS3 | CCDC33 |
| GMCL1P1 | COMMD6 | SPATA31E1 | ABRACL | SAMD11 | UBQLN4 |
| TCEA3 | SYT2 | INSL4 | RAG1 | CCNL2 | CRP |
| DDX11L1 | LOC729737 | WASH7P | LOC100132287 |  |  |

**Table S12.**
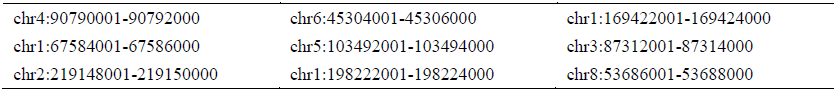
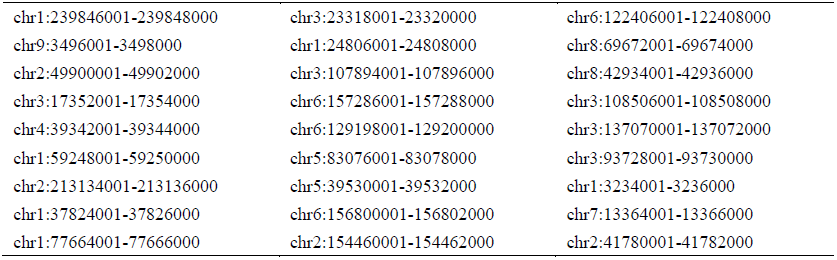
Top DhMR feature set used for cancer prediction.

|  |  |  |
| --- | --- | --- |
| chr4:90790001-90792000 | chr6:45304001-45306000 | chr1:169422001-169424000 |
| chr1:67584001-67586000 | chr5:103492001-103494000 | chr3:87312001-87314000 |
| chr2:219148001-219150000 | chr1:198222001-198224000 | chr8:53686001-53688000 |
| chr1:239846001-239848000 | chr3:23318001-23320000 | chr6:122406001-122408000 |
| chr9:3496001-3498000 | chr1:24806001-24808000 | chr8:69672001-69674000 |
| chr2:49900001-49902000 | chr3:107894001-107896000 | chr8:42934001-42936000 |
| chr3:17352001-17354000 | chr6:157286001-157288000 | chr3:108506001-108508000 |
| chr4:39342001-39344000 | chr6:129198001-129200000 | chr3:137070001-137072000 |
| chr1:59248001-59250000 | chr5:83076001-83078000 | chr3:93728001-93730000 |
| chr2:213134001-213136000 | chr5:39530001-39532000 | chr1:3234001-3236000 |
| chr1:37824001-37826000 | chr6:156800001-156802000 | chr7:13364001-13366000 |
| chr1:77664001-77666000 | chr2:154460001-154462000 | chr2:41780001-41782000 |

